# SCPattern: A statistical approach to identify and classify expression changes in single cell RNA-seq experiments with ordered conditions

**DOI:** 10.1101/046110

**Authors:** Ning Leng, Li-Fang Chu, Jeea Choi, Christina Kendziorski, James A. Thomson, Ron Stewart

## Abstract

**Motivation:** With the development of single cell RNA-seq (scRNA-seq) technology, scRNA-seq experiments with ordered conditions (e.g. time-course) are becoming common. Methods developed for analyzing ordered bulk RNA-seq experiments are not applicable to scRNA-seq, since their distributional assumptions are often violated by additional heterogeneities prevalent in scRNA-seq. Here we present SC-Pattern - an empirical Bayes model to characterize genes with expression changes in ordered scRNA-seq experiments. SCPattern utilizes the non-parametrical Kolmogorov-Smirnov statistic, thus it has the flexibility to identify genes with a wide variety of types of changes. Additionally, the Bayes framework allows SCPattern to classify genes into expression patterns with probability estimates.

**Results:** Simulation results show that SCPattern is well powered for identifying genes with expression changes while the false discovery rate is well controlled. SCPattern is also able to accurately classify these dynamic genes into directional expression patterns. Applied to a scRNA-seq time course dataset studying human embryonic cell differentiation, SCPattern detected a group of important genes that are involved in mesendoderm and definitive endoderm cell fate decisions, positional patterning, and cell cycle.

**Availability and Implementation:** The SCPattern is implemented as an R package along with a user-friendly graphical interface, which are available at:https://github.com/lengning/SCPattern

## 1 Introduction

RNA sequencing (RNA-seq) technology has been replacing microarray based gene expression methods because of its improved accuracy and reduced cost per sample. RNA-seq of a population of cells in bulk (bulk RNA-seq) has been used in a number of experiments to characterize expression differences between biological conditions. Among these experiments, ordered bulk RNA-seq experiments are becoming quite popular. Ordered bulk RNA-seq experiments measure changes over multiple ordered conditions (e.g. over time or space). Instead of classifying genes as differentially expressed (DE) or equally expressed (EE) as in an experiment comparing two conditions, the ordered experiments provide precise views of graded changes.

Recent advances in single cell technologies allow investigators to use RNA-seq technology to characterize transcriptome-wide expression measures of individual cells. Compare to ordered bulk RNA-seq experiments which measure averaged expression over thousands of cells for a given time point or position, ordered scRNA-seq experiments provide unprecedented power to understand how a cell population’s expression distribution changes during a biological process. Recent ordered scRNA-seq studies include experiments studying differentiation of human T cells (Buettner *et al*., 2015), human myoblasts (Trapnell *et al*., 2014), and human preimplantation development (Yan *et al*., 2013; Blakeley *et al*., 2015; Töhönen *et al*., 2015), to name a few.

In experiments with ordered conditions, investigators are mainly interested in identifying genes with expression changes along the multiple conditions (DE genes) and classifying them by their expression patterns (e.g. Up-Up-Up-Up, Up-Up-Down-Down, etc.). A number of methods have been developed for DE gene identification and pattern specification for ordered bulk RNA-seq experiments (Nueda *et al*., 2014; Leng *et al*., 2015). However, these methods are not directly applicable to scRNA-seq data, because they typically assume that the gene expression follows a unimodal distribution within each condition. This assumption is not true in scRNA-seq data due to cell heterogeneity and to technical dropouts (cells having zero counts due to technical reasons).

In a two-condition scRNA-seq experiment, investigators typically treat zero counts as technical dropout events. This is because without additional information, it is impossible to determine if the zero counts are biological or technical in basis. For example, a Bayesian model has been developed to detect DE genes in a two-condition scRNA-seq experiment (SCDE (Kharchenko *et al*., 2014)). The SCDE model uses a mixing parameter to account for the dropout events, and a location parameter to estimate the mean expression level of expressed cells. In the SCDE implementation, a statistical test is provided to detect mean differences of expressed cells, but no guideline is provided to characterize differences due to percentage of zero changes, and extra heterogeneity among expressed cells that gives rise to distributions that are not unimodal is not accommodated.

Unlike two-condition experiments, in an ordered scRNA-seq experiment, investigators may infer whether the zero counts are mainly due to biological or technical sources by incorporating information from other time points (positions). For example, gradual changes in the percentage of zeros over ordered conditions is more consistent with a biological signal, compared with the case that percentages of zeros remain relatively constant. Therefore, it is important to detect both mean expression changes and percentage of zero changes in an ordered scRNA-seq experiment. Figure 1(a-c) shows examples from three classes of DE genes in an scRNA-seq time course experiment - genes whose mean expression of non-zero cells changes over time; genes whose percentage of zero cells changes over time; and genes with both changes. Only considering mean expression changes in non-zero cells will ignore genes in the same class as those in Figure 1(b), resulting in reduced power for identifying meaningful changes across conditions. In addition, currently there is no scRNA-seq method that allows for directional tests, and consequently it is not clear how to classify genes into patterns in an ordered scRNA-seq experiment. To address these challenges, we developed an empirical Bayes approach SCPattern which identifies genes with expression changes by considering cells with zero and non-zero cells collectively, and classifies them into directional expression patterns with probability estimates. A graphical user interface (GUI) implementation is also available.

**Figure 1:**
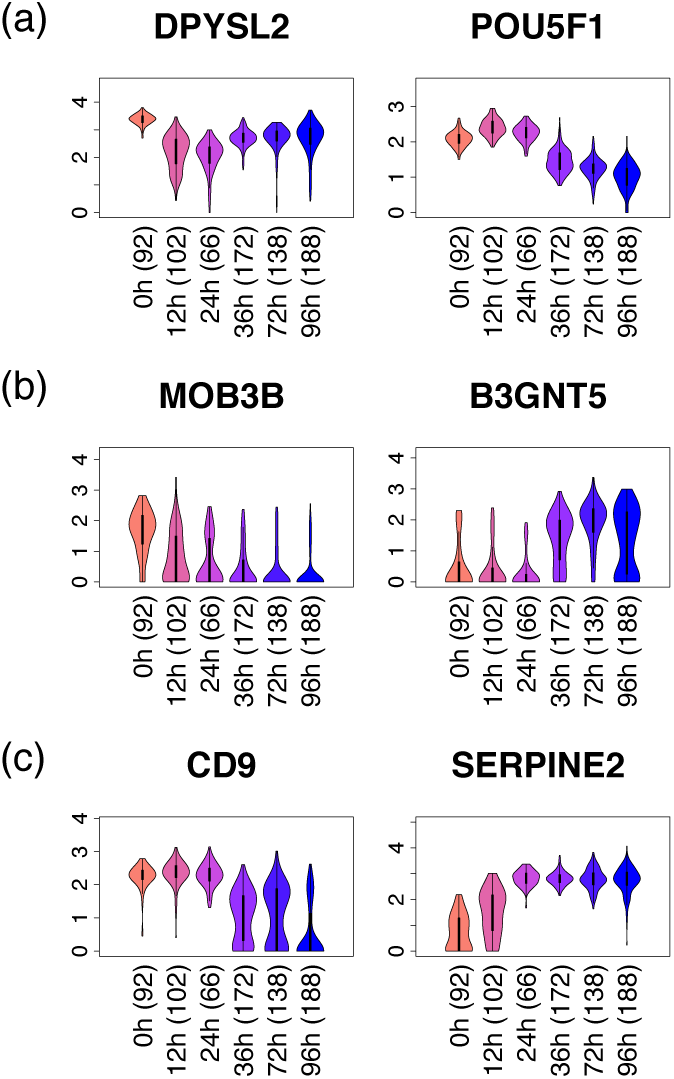
Example genes in the scRNA-seq time course data studying definitive endoderm cell differentiation. Panels (a), (b) and (c) show example genes with expression mean changes in non-zero cells, percentage of zero changes and both, respectively. The x-axis shows time points and the y-axis shows log10 (normalized expression + 1). One was added to avoid showing negative infinity values.

Simulation studies suggest that considering non-zero counts only in analysis of ordered scRNA-seq experiments results in reduced power for detecting DE genes. On the other hand, SCPattern provides improved power while its false discovery rate (FDR) is well controlled. In addition to DE gene detection, results also show that SCPattern has high accuracy in classifying genes into expression patterns. We also compared the algorithms with a naive method based on fold change (FC), which was used as a screening criteria in many scRNA-seq studies (Yan *et al*., 2013; Blakeley *et al*., 2015; Finak *et al*., 2015). Results show that the FC method’s operating characteristics are sensitive to the choice of cutoff, and its FDR is always inflated. Similar results are demonstrated in a case study of a time course experiment of differentiating human embryonic stem cells (ES cells) towards definitive endoderm cells (Chu* *et al*., 2015).

## 2 Methods

### The SCPattern algorithm

To simplify the presentation, we refer to ordered levels as time points, noting that SCPattern accommodates other ordered data structures such as ordered in space, by gradient, etc. Denote the time points by *t* = 1,2,…, *T*. Let 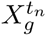 be expression of gene *g*, cell *n* from time *t*. Then 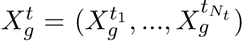 denotes gene *g*’s expression at time point *t*, in which *N*_*t*_ is the number of cells in time point *t*.

Comparing gene expressions between time *t* and *t* + 1, SCPattern tests whether a gene’s distribution changes across these two conditions. To characterize the distributional differences, SCPattern utilizes the directional Kolmogorov-Smirnov (K-S) statistic. Given a value *ω*, denote the cumulative distribution functions (CDFs) of 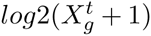 and 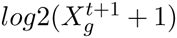 by 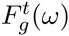 and 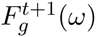, respectively (one was added to each value to avoid generating negative infinity values and to reduce dynamic range of low expressed genes). Then the K-S statistics for up‐ and down-regulation are:

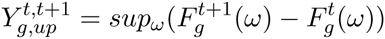

and

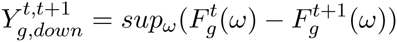

We use the K-S statistic to characterize expression differences because it accommodates both expression level changes and percentage zero changes.

Examples are shown in Figure 2. Specifically, 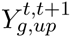 is large when gene *g*’s non-zero cells have higher expression at time *t* + 1, when more cells become non-zero at time *t* + 1, or both (Figure 2(a)). Similarly, 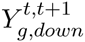 is large when gene *g*’s non-zero cells’ expression level decreases, when more zero cells present at time *t* + 1, or both (Figure 2(b)). Both 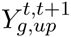 and 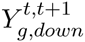 are small when there is no expression change between the two time points (Figure 2(c)).

**Figure 2:**
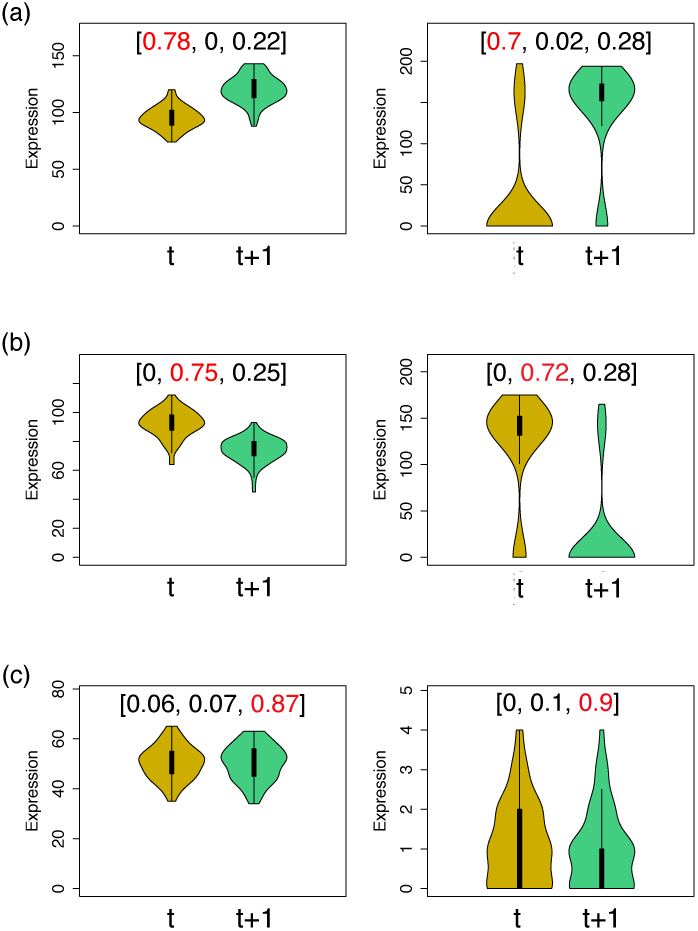
Example genes and their summary statistics in a comparison between two time points. Denote the two time points as *t* and *t* + 1. Panels (a), (b) and (c) show genes that are up-regulated, down-regulated and EE, respectively. The y axis indicates gene expression. The vector 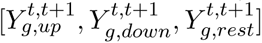 is shown in the brackets for each gene. The largest value of each gene is marked as red.

For each pair of time points, by taking genes with large 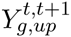, genes with large 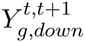, and genes with large 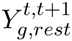, we can classify genes into categories Up, Down and EE, respectively. Here 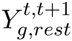 is defined as 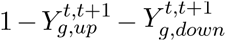. Note that since a CDF is a monotonic function, 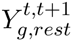 is always greater or equal to zero. Denote 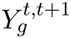 as a vector 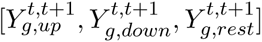.

Since 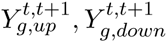 and 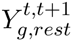 all lie between 0 and 1, 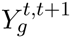 is well described by a Dirichlet distribution. To classify genes into these three categories, we assume 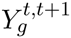 follows a mixture of Dirichlet distributions:

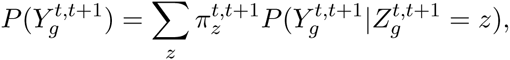

in which *z* is a latent variable indicating Up, Down or EE. The conditional probability 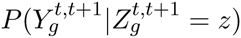 is defined as

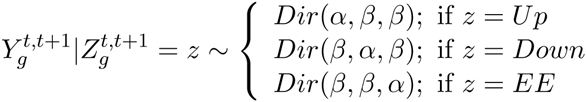

with constraint *α* > *β*. Note that for a Dirichlet distribution Dir(*γ*_1_, *γ*_2_, *γ*_3_), the probability density function is defined as

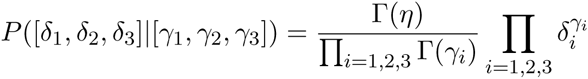

where *η* = *γ*_1_+*γ*_2_+*γ*_3_; and its expected values are defined as (*γ*_1_/*η*, *γ*_2_/*η*, *γ*_3_/*η*). In our parameterization, 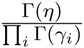 remains the same for all three Dirichlet components. Therefore, for any *z*0, *z*1 in Up, Down and EE, our model ensures that when 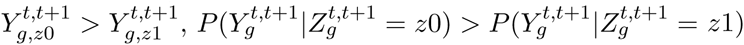. For example, in a case when 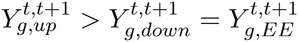, to compare the conditional probabilities across three components we have:

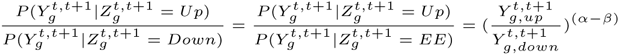

which is always greater than 1.

Thus, in a time course experiment with *T* time points, the marginal distribution along all time points can then be written as:

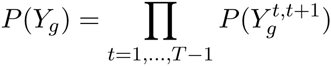

By Bayes rule, the posterior probability (PP) of being pattern *k*_1_ is then calculated as

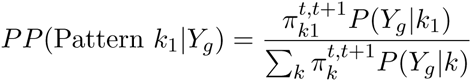

For example, suppose *k*_*m*_ represents pattern Up-Up-Up in a time course with 4 time points, then

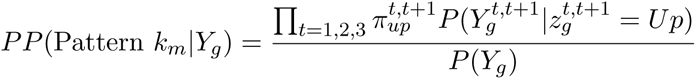

To estimate parameters *α* and *β*, SCPattern generates *B* permuted genes by randomly shuffling the cells across the conditions. The resulting permuted genes are expected to be EE. Therefore, SCPattern estimates *α* and *β* by assumin 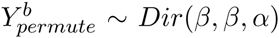 for all *B* permuted genes. *B* defaults to 100000 and can be specified by the user. Once *α* and *β* are estimated, the mixing parameter *π*s are estimated via Expectation-Maximization algorithm and the PPs can then be obtained. A gene’s mostly likely pattern (MLP) is then defined as the pattern that has the highest PP among all possible patterns. A gene’s MLP is also its maximum a posteriori estimate.

### User-friendly graphical interface

SCPattern is available as an R package (R/SCPattern) along with a user-friendly GUI. It takes as input estimates of gene expression and a vector that specifies the condition to which each cell belongs. The output of the SCPattern GUI contains a list of DE genes and their PPs, the most likely pattern of each DE gene, and violin plots. Figure 3 shows the SCPattern-GUI interface.

**Figure 3:**
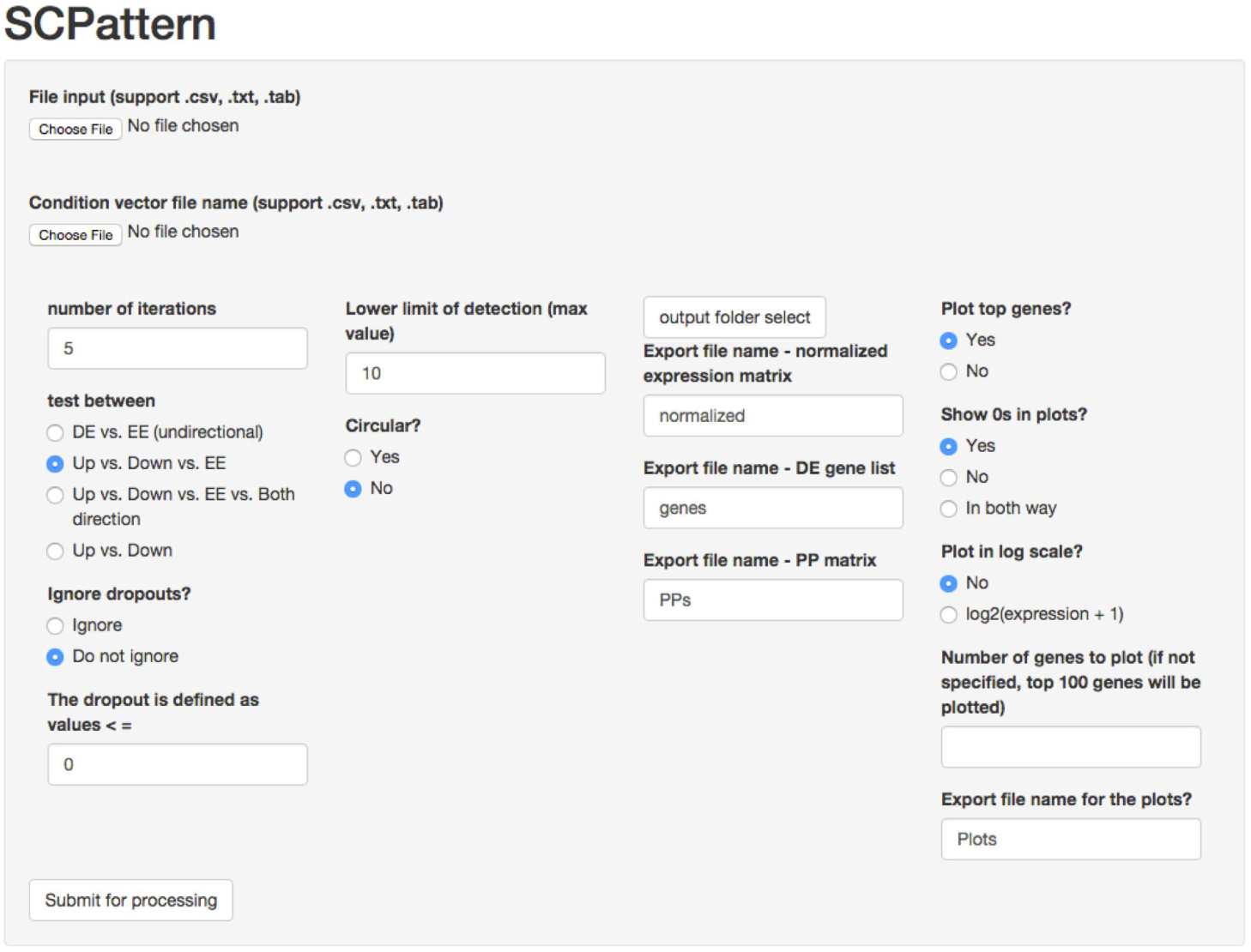
Screenshot of the SCPattern-GUI. The SCPattern-GUI is implemented using R/shiny package.

### Comparison to other methods on simulated data

We followed a typical simulation setup (Robinson and Smyth, 2007; Leng *et al*., 2013, 2015) by generating counts from a Negative Binomial distribution. The gene-specific mean and variance of time point *t* are defined as *μ*_*gt*_ and *μ*_*gt*_(1 + *μ*_*gt*_*ϕ*_*gt*_), respectively. The (*μ*_*gt*_, *ϕ*_*gt*_) values were sampled as pairs from the case study data in the definitive endoderm cell differentiation experiment (See Supplementary Note section 2 for more details). FC for DE genes were sampled from empirical FCs calculated using empirical data. Each simulated dataset contains 10000 genes. We generated 100 datasets for each simulation scenario.

**Sim I** was conducted to benchmark SCPattern in comparison between two adjacent time points *t* and *t* + 1. Sixty cells were simulated for each of the two time points. We simulated 3000 of the 10000 genes to be Up or Down from *t* to *t* + 1. In addition, 1000 of these 3000 genes were simulated to have changes in the percentage of cells having zero expression. The other 2000 genes were simulated to be non-zero and have mean change across the two time points. Among all 10000 genes, we simulated 2000 to have a bi-model expression distribution for the non-zero cells in at least one time point. Technical dropouts (stochastic zero counts due to technical reasons) were also simulated. For any gene, we randomly selected 10% of the cells and simulated them as technical dropouts by setting their expression value to 0.

**Sim II** was conducted to evaluate SCPattern and other methods in a time course experiment. In each simulation we simulated 5 time points, each with 60 cells. We simulated 3000 genes to be changing over time - 1000 have changes in the percentage of cells having zero expression and the other 2000 are non-zero and have mean changes. We simulated 2000 genes to have a bi-modal expression distribution in at least one of the time points. The technical dropouts were simulated as in **Sim I**.

To identify a list of DE genes using SCPattern, we take those genes whose MLP is not the constant pattern. The constant pattern is defined as “EE” in **Sim I** and “EE-EE-EE-EE” in **Sim II**. Recall that SCPattern provides a gene-specific PP associated with each expression pattern. The MLP of gene *g* is then defined as argmax_*k*_ (*PP*(*Pattern*_*k*_)). In **Sim II**, after calling a gene as DE, SCPattern classifies the gene into its MLP.

We included SCDE in **Sim I** evaluations. A gene is called DE by SCDE if its p-value is less than 0.05. The p-values were calculated from z-scores provided by the SCDE package. Since SCDE was not designed to classify genes into directional expression patterns in time course data (and no information is provided in their user manual on how to do so), it is not included in the evaluations of **Sim II**. To mimic SCDE and illustrate how treating all zero counts as technical dropouts affects analysis results, we also implemented SCPattern in a way that only considers the non-zero values (SCPattern-nonzero). SCPattern-nonzero is implemented in a similar way to SCPattern. In SCPattern-nonzero, for each gene, prior to further analysis, we removed cells whose expression value is zero. Therefore, the K-S statistics are calculated using only the non-zero values in a gene. Similarly, the permutations are also conducted using the non-zero values exclusively. SCPattern-nonzero was included in all simulation studies.

We also evaluated an implementation of naive FC methods. For the naive FC method, denote 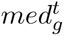 as the median expression of gene *g* at time point *t*. A gene *g* is called Up (Down) between *t* and *t* + 1 if 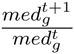 is greater than (less than) *m*; otherwise, it is called EE. We considered three values of *m*: 1.2, 1.5, and 2. In **Sim II**, a gene is defined as DE if it is Up or Down in any pair-wise comparison.

When evaluating different methods’ performance to detect DE genes, the power and false discovery rates were calculated as:

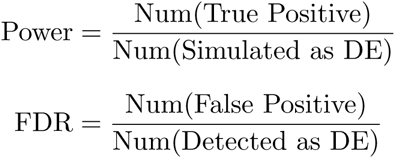

When evaluating the performance in classifying genes into patterns, we consider:

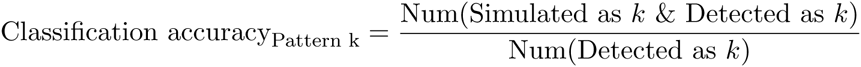

## 3 Results

### 3.1 Simulation studies

As described above, two simulation studies were conducted to investigate the operating characteristics of SCPattern, SCDE, SCPattern-nonzero and naive FC methods. Table 1 shows the power and FDR for identifying DE genes in **Sim I**. In addition to showing the overall power, we also evaluated the statistical power within gene subgroups with non-zero mean change and percentage zero change (FDR is not shown for these subgroups because false discoveries cannot be classified into non-zero mean change or percentage zero change groups). As shown in Table 1, SCPattern has higher overall power and better controlled FDR than SCDE and SCPattern-nonzero. This is largely because SCDE and SCPattern-nonzero lack power in identifying DE genes with percentage zero changes. On the other hand, all three methods have reasonable power in identifying genes with non-zero mean changes. As expected, the power and FDR of the naive FC method is very sensitive to the choice of the cutoff *m*. When *m* = 1.2, the naive FC method has highest overall power among all methods, but an inflated FDR (16.3%). The power of the naive FC method drops dramatically as *m* increases. In addition, in empirical data analyses where underlying truth is unknown, it is not clear how to pick an appropriate threshold.

**Table 1:**
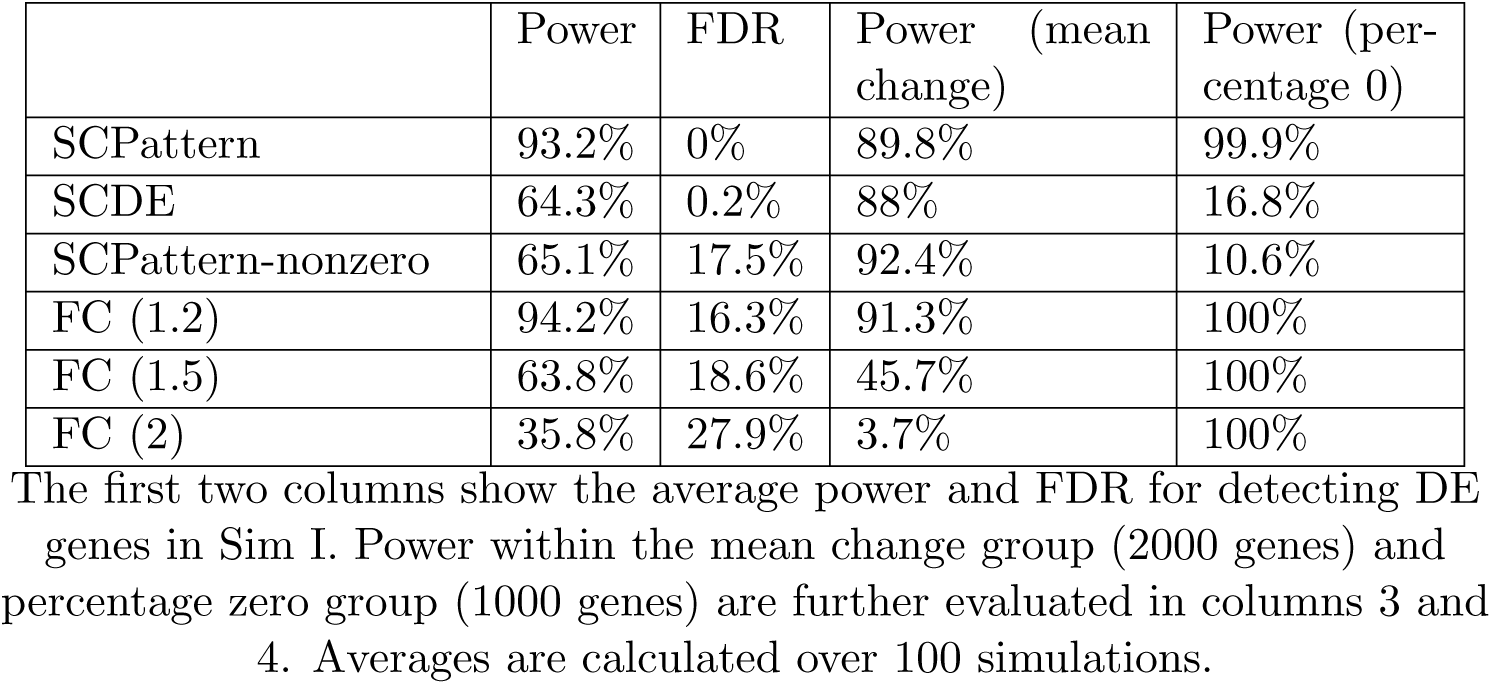
Operating characteristics for identifying DE genes in **Sim I**

Table 2 shows power and FDR for identifying DE genes in **Sim II**. Similar to **Sim I**, SCPattern outperforms SCPattern-nonzero because of its improved power in identifying genes with percentage zero changes. Compare to other methods, the naive FC method has high FDR in all cases. We also evaluated SCPattern in classifying DE genes into expression patterns. Figure 4 shows the classification accuracy for the top 10 patterns simulated in **Sim II**. As detailed above, the classification accuracy of a pattern *k* is defined as the percentage of correct classification among genes that were classified into pattern *k*. Results in Figure 4 indicate that SCPattern successfully classified most of the genes into the correct pattern.

**Table 2:**
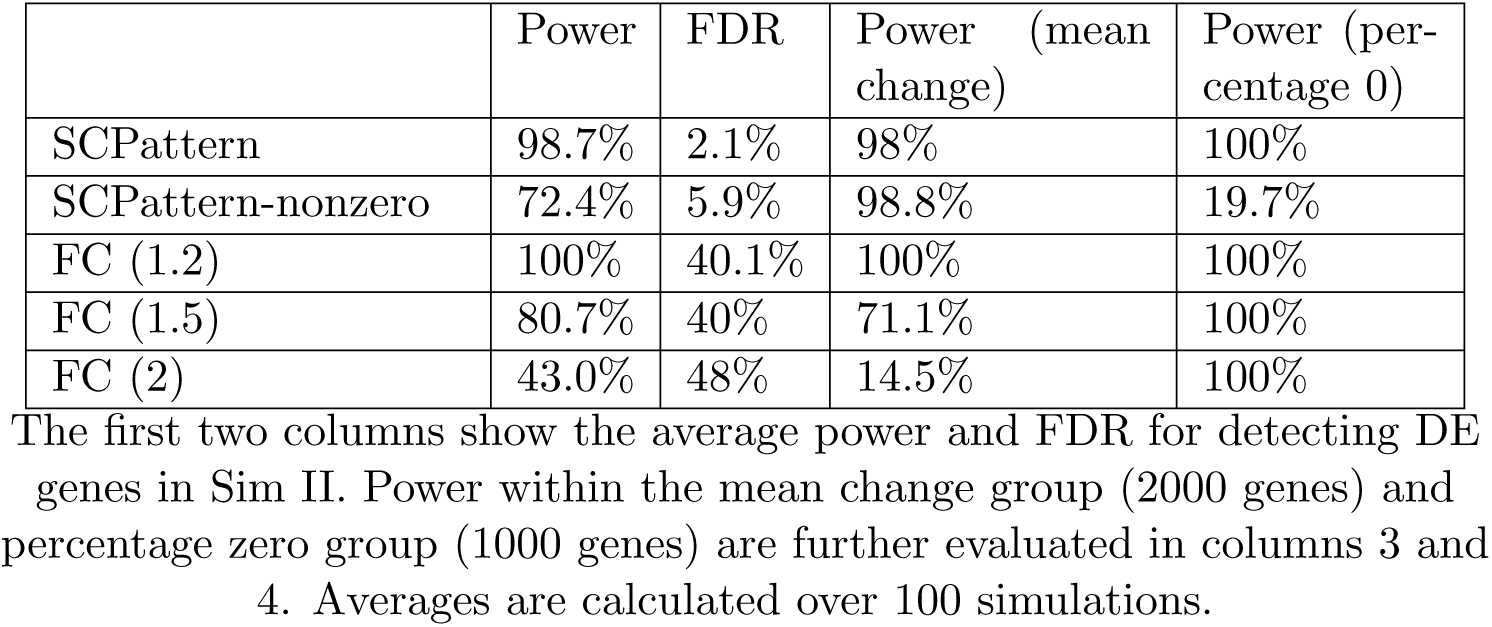
Operating characteristics for identifying DE genes in **Sim II**

**Figure 4:**
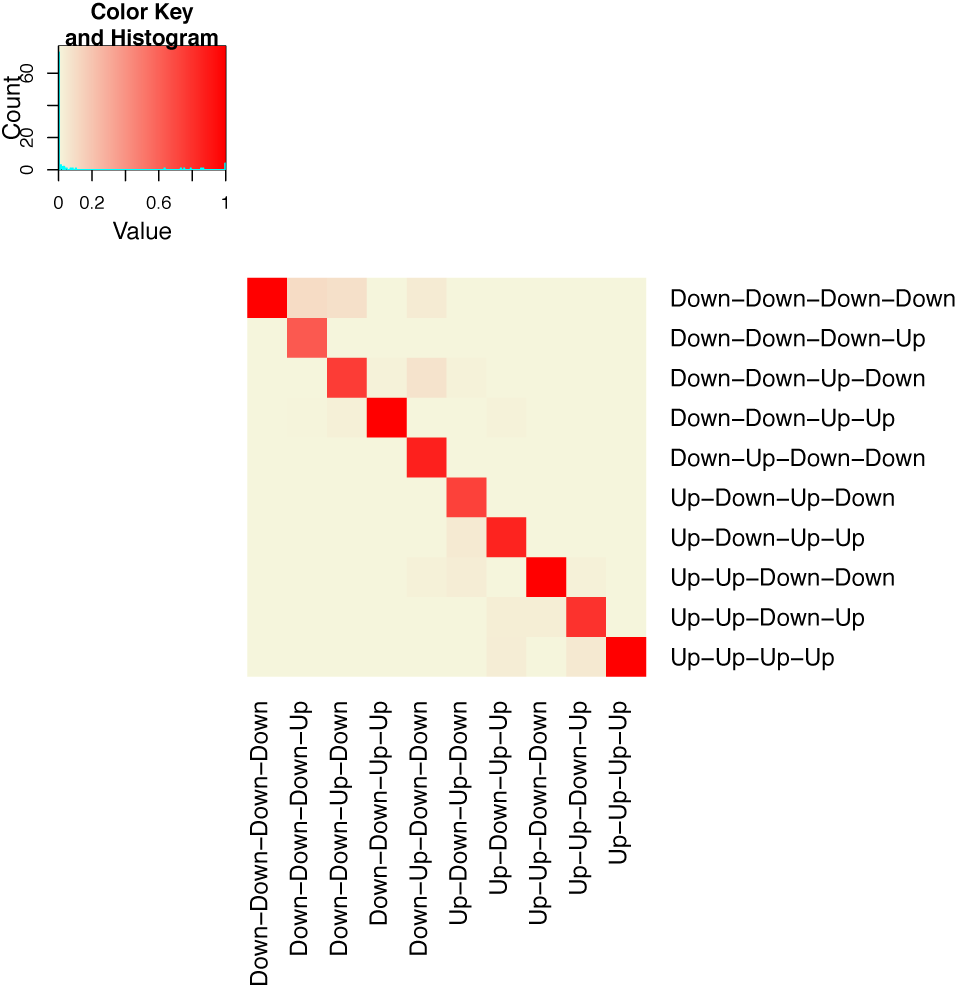
The classification accuracy of the top 10 expression patterns in Sim II. Rows represent the simulated patterns and columns represent classified patterns determined by SCPattern. The color in pixel (*i*, *j*) shows the fraction of genes that were simulated as pattern *i* and classified as pattern *j*. The results shown are averages from 100 simulations.

### SCPattern identifies definitive endoderm differentiation markers in the human ES cell scRNA-seq dataset

Of interest in our case study, detailed below, is scRNA-seq data from a differentiation time course experiment from human ES cells to definitive endoderm cells over a 4 days period (Chu* *et al*., 2015). Six time points were considered in this time-course experiment. A total of 758 cells were sequenced, in which 92, 102, 66, 172, 138 and 188 cells were considered for analysis at 0h, 12h, 24h, 36h, 72h and 96h, respectively.

We applied SCPattern, SCPattern-nonzero, and naive FC methods on the scRNA-seq time course data set. A total of 3247, 4581, 9327, 10074 and 10301 genes were identified by SCPattern, SCPattern-nonzero, and FC method with target FC 2, 1.5 and 1.2, respectively (Supplementary Table 1). Among those genes identified by SCPattern, 883 were not detected by SCPattern-nonzero (which is designed to mimic existing two condition DE methods such as SCDE if they could be applied to ordered data). As in the simulation study, the majority of them are DE genes with percentage zero changes. Figure 5(a) shows expression violin plots for 5 of these genes. In Figure 5(a), the left(right) column shows plots that contain(exclude) the zero counts. All these genes have constant expression levels for the non-zero cells but the percentage of zero cells changes gradually. Excluding zero cells in the analysis results in missing these changing genes.

**Figure 5:**
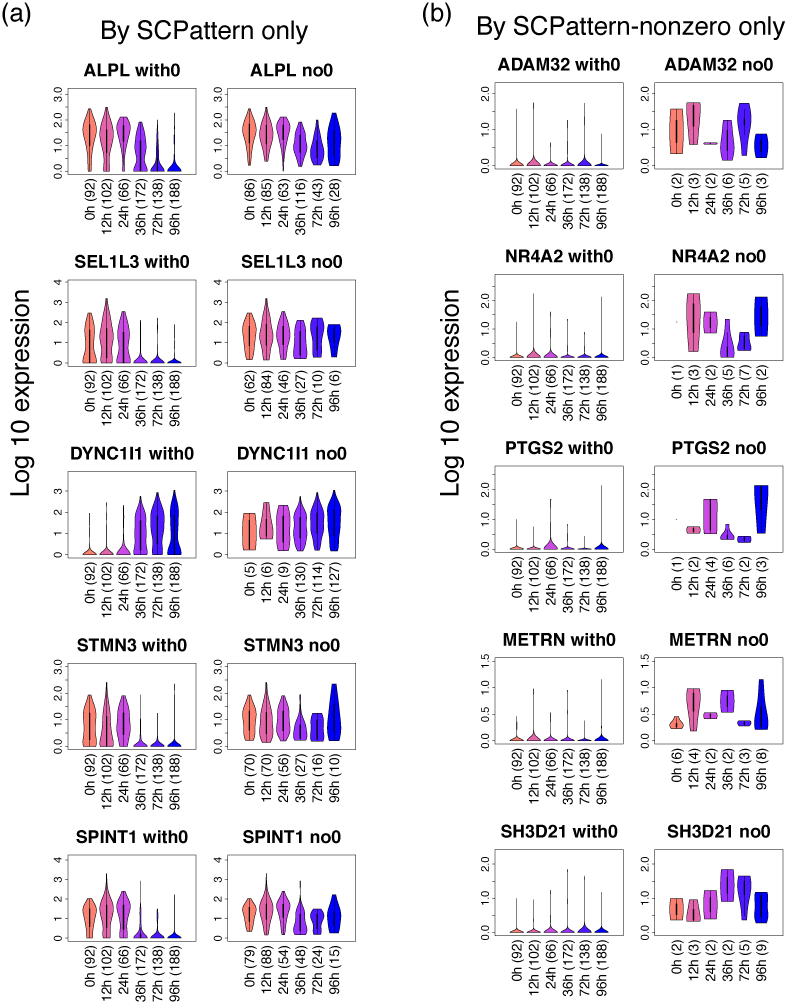
Example genes that were exclusively identified by SCPattern or SCPattern-nonzero. Panel (a) shows example genes identified by SCPattern but not SCPattern-nonzero in the scRNA-seq data studying definitive endoderm cell differentiation. The left column shows violin plots of log10 (normalized expression +1). Cells from different time points are shown in different colors. The right column is similar to the left column, but only show cells with non-zero expression within a gene. Panel (b) shows example genes identified by SCPattern-nonzero but not SCPattern in the case study data.

Among the 883 genes exclusively identified by SCPattern, a group of of them are known to be important to definitive endoderm differentiation. Figure 6(a) shows eight example genes that were identified by SCPattern but not SCPattern-nonzero. Among these eight genes, *MIXL1*, *DKK1*, *EOMES* and *PITX2* show an up-regulated trend at the beginning of the differentiation. Each of these 4 genes is known to be involved in mesendoderm or definitive endoderm cell fate decisions (Faucourt *et al*., 2001; Costello *et al*., 2011; Lim *et al*., 2009; Lewis *et al*., 2007). In addition to these four genes, Figure 6(a) shows two *HOX* genes that follows the EE-EE-EE-Up-EE pattern. The *HOX* gene families are known for the establishment of anterior-posterior patterning along the body axis (Guazzi *et al*., 1998; Zakany and Duboule, 2007). Furthermore, we found a group of cell cycle genes with a down-regulated trend at 72h. In particular, cell cycle regulators *MYCN* and *PLK1* were detected as EE-EE-EE-Down-EE (Figure 6(a)). This indicates a change of cell-cycle regulation during definitive endoderm cell differentiation (Tan *et al*., 2013). Additional classification results may be found in Supplementary Note section 3.

**Figure 6:**
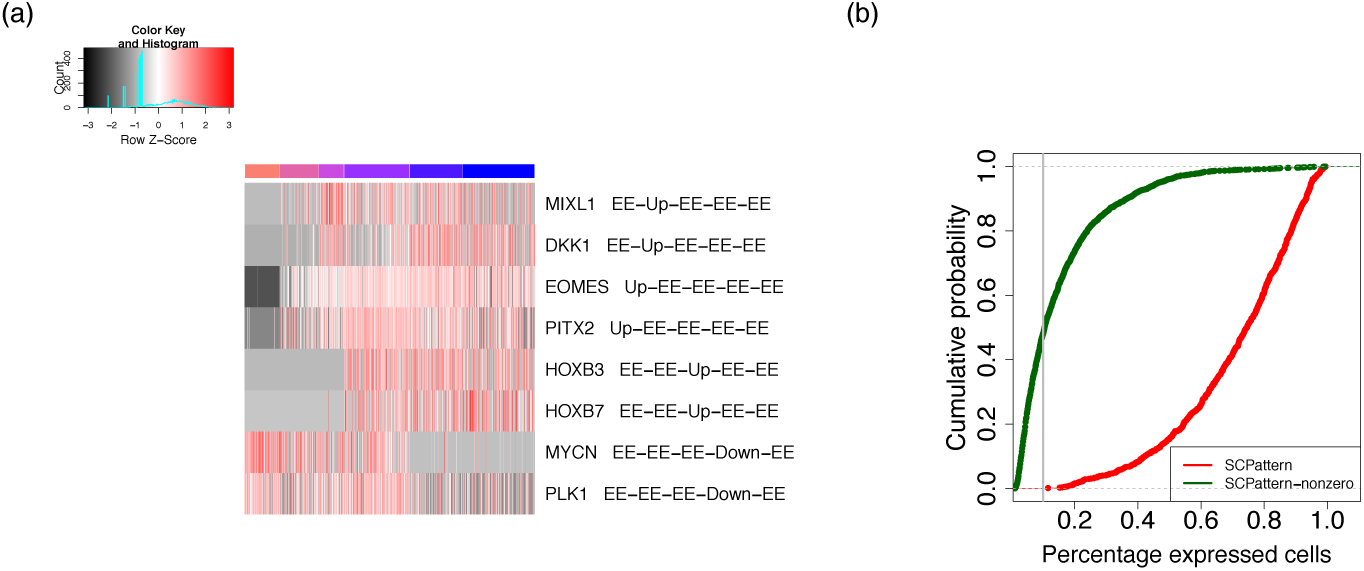
Case study results comparing SCPattern and SCPattern-nonzero. Panel (a) shows eight example genes that were identified by SCPattern but not SCPattern-nonzero. Each row represents one gene and the associated MLP is also included in the row name. The gene expression values were rescaled to gene-specific z-scores. The columns show cells following time course order. Within each time point the cells were sorted by their sample ID. Cells from different time points are marked in different colors. Panel (b) shows the CDF of percentage expressed cells in genes exclusively identified by SCPattern and SCPattern-nonzero, respectively.

Also of interest are 2217 genes that were identified by SCPattern-nonzero but not SCPattern. We found that most of these genes have very few expressed cells. Five example genes among these 2217 are shown in Figure 5(b). Although the expression level of non-zero cells changes over time, the vast majority of the cells are not expressed. And though these changes in the minority of expressed cells might represent biological signals, it would be challenging to perform downstream biological validation on genes with few expressed cells.

To further investigate the characteristics of the 883 genes exclusively identified by SCPattern vs. the 2217 genes exclusively identified by SCPattern-nonzero, we evaluated the percentage of expressed cells for these genes. Figure 6(b) shows the empirical CDFs of the percentage of expressed cells for these two groups of genes. The CDFs indicate that nearly half of the 2217 genes exclusively identified by SCPattern-nonzero are expressed in less than 10% of the cells. Among the 883 genes exclusively identified by SCPattern, all of them are expressed in more than 10% of the cells, and over 80% are expressed in more than half of the cells. Taken together, these results suggestthat SCPattern-nonzero is likely to identify genes who contain a large proportion of zero cells.

## 4 Discussion

We have developed SCPattern - a statistical method for characterizing genes having expression changes in an ordered scRNA-seq experiment. SCPattern is an empirical Bayes model that utilizes the K-S statistic, which does not rely on any parametrical assumptions. Therefore, it is not limited with respect to the types of expression changes that may be detected. Once DE genes are identified, SCPattern further classifies them into directional expression patterns. Additional down-stream analyses of DE genes may then be conducted to characterize sub-populations within a condition (Feigelman *et al*., 2014), or to recover the progression trajectory (Trapnell *et al*., 2014; Shin *et al*., 2015).

The SCPattern method is implemented as an R package (R/SCPattern).

We also provide an implementation of a user friendly GUI, which allows users with little computational knowledge to perform analyses. Simulation and case study results show that SCPattern provides improved power over SCPattern-nonzero which ignores zero counts in the analyses of ordered scRNA-seq experiments. This is mainly because SCPattern has higher power in identifying genes with percentage zero changes along the ordered conditions. For studies that exclusively focus on changes in expressed cells, the SCPattern-nonzero implementation is also available in the R/SCPattern package and its GUI. Additional investigation of the SCPattern case study results is detailed in a companion study (Chu* *et al*., 2015). On the other side, results also show that the FC method’s performance is very sensitive to the choice of cutoff, while it is challenging to determine the optimal FC cutoff in empirical data analysis. Thus, extra cautions might be needed when detecting DE genes by purely FC thresholding or by combining statistical tests with FC thresholding (Blakeley *et al*., 2015; Yan *et al*., 2013; Finak *et al*., 2015).

In this manuscript, we present SCPattern results to classify each transition into categories Up, Down and EE. However, the SCPattern empirical Bayes model could be generalized to accommodate other categories of interest. The R/SCPattern package provides several additional options to define the categories. For example, it is also possible to incorporate a fourth category that gene expression distribution changes in both directions (some cells have increased expression from *t* to *t* + 1 while the others decrease, see Supplementary Figure 1 for an example). Although we have rarely seen this type of change in our case study data, they might be of interest in other studies. More details may be found in Supplementary Note section 1.

## Acknowledgements

This work was supported by the National Institutes of Health 4UH3TR000506-03, 5U01HL099773-06, GM102756, U54 AI117924, the Charlotte Geyer Foundation, and the Morgridge Institute for Research. We thank J. Bolin, A. Elwell, and B.K. Nguyen for the preparation and sequencing of the RNA-seq samples. We thank P. Jiang and S. Swanson for performing the RNA-seq read processing.

